# Profiles and Dynamics of the Transcriptome of Microglial Cells Reveal their Inflammatory Status

**DOI:** 10.1101/2021.12.20.473548

**Authors:** Keren Zohar, Elyad Lezmi, Fanny Reichert, Tsiona Eliyahu, Shlomo Rotshenker, Marta Weinstock, Michal Linial

## Abstract

The primary role of microglia in the maintenance of brain homeostasis is to respond to disturbances in the microenvironment. In this study, we cultured murine neonatal microglia and activated them with lipopolysaccharide (LPS) and benzoyl ATP (bzATP) to characterize changes in the transcriptome in response to various *in vivo* stimuli caused by pathogens, injury, or toxins. Activation by bzATP, an agonist of purinergic receptors, induces a transient wave of transcriptional changes. However, a long-lasting transcriptional profile affecting thousands of genes occurs only following a combination of bzATP and LPS. This profile is dominated by a coordinated induction of cytokines (e.g., IL1-α and IL1-β), chemokines, and their direct regulators. Many of these inflammatory-related genes are up-regulated by several orders of magnitude. We identified the TNF-α and NF-κB pathways as the principal hubs for signaling of interleukin and chemokine induction in this cell system. We propose that primary microglia under controlled activation paradigms can be used for testing reagents that could attenuate their activated state. Such a microglial system could serve as a model for changes occurring in the aging brain and neurodegenerative diseases.

**Highlight points:**

* Primary murine microglia cultures release cytokines following activation with bzATP and LPS
* The wave of changes in gene expression by bzATP is transient.
* bzATP+LPS causes a transcription program dominated by the induction of interleukins and chemokines.

## Introduction

Microglia act as the resident macrophages of the central nervous system (CNS). Their function is to maintain brain homeostasis and respond effectively to a broad spectrum of perturbations induced by acute stress like toxic agents or physical injury [1-3]. This is accomplished through communication with the surrounding neurons and astrocytes and the release of signaling molecules like ATP (adenosine tri-phosphate), that interact with specific receptors on the microglial membrane [4, 5]. To protect neurons from apoptotic death and remove dying cells, microglia respond by changing their morphology and function [6-8]. They release cytokines, reactive oxygen, and nitrogen species (ROS and RNS, respectively, like nitric oxide (NO) [9, 10]. During chronic stress and neuronal damage that occurs during aging, the release of pro-inflammatory cytokines becomes prolonged and together with oxidative stress, exacerbates neurodegeneration [11]. Thus, glial cells with excess amounts of neuroinflammatory markers like HLA-DR, CD68, and CD105 are abundant in Parkinson’s (PD), Alzheimer’s diseases (AD) and frontotemporal dementia [12].

To better understand the processes involved in the regulation of microglial activity, studies have been performed under controlled conditions in isolated microglia, prepared from neonatal rodents and cultured alone or with astrocytes [13]. Exposure of these cells to activators like lipopolysaccharide (LPS), interferon-gamma (IFNγ), interleukins, and other extracellular signaling molecules also produces a change in their function, morphology and inflammatory responses [14, 15]. In microglia, LPS increases the release of TNF-α and other pro-inflammatory cytokines by activating the PI3K and AKT pathway and induces the nuclear translocation of NF-κB [9, 16, 17].

RNA sequencing analysis (RNA-seq), used for quantifying gene expression, has revealed the contribution of microglia to inflammation following injury [18] and their mode of activation during the aging process [19, 20]. In this study, we used RNA-seq methodology in cultured, neonatal microglia to obtain a molecular view of the dynamics of the transcriptome and cytokine release in response to activation by LPS with and without the addition of benzoyl ATP (bzATP), an agonist of the P2RX7 subtype of ATP receptors.

## Materials and Methods

### Compounds and reagents

Dulbecco’s Modified Eagle Medium (DMEM), DMEM/F12, Gentamycin sulfate and L-Glutamine were obtained from Biological Industries (Beit-Haemek, Israel), and 2’-3’-O-(4-benzoyl benzoyl) adenosine 5’-triphosphate (bzATP), Bovine serum albumin (BSA) and LPS were purchased from Sigma-Aldrich (Israel).

### Preparation of microglial cultures

Primary microglia were isolated from the brains of neonatal male Balb/C mice (Harlan Sprague Dawley, Inc., Israel) as described in [21]. The cells were plated in Poly-L-lysine coated flasks for one week and all non-adherent cells washed away by re-plating for 1-2 hours (hrs) on bacteriological plates. Microglial cells were propagated by supplementing the culture with 20% of medium conditioned from L-cells that produce mouse-CSF (colony-stimulating factor). In all experiments the heat-inactivated fetal calf serum (FCS) was replaced by purified BSA. The identity of the microglia was validated by their morphology and by several molecular markers using immunostaining by P2Y1, F4/80, CR3 (complement receptor-3), and Galectin-3/MAC-2 [22].

### Measurement of cytokines

The activation of the cultured microglia was measured by cytokine release as previously described [23] and according to manufacturer’s protocols. Cells were grown to 75% confluence in 6-well plates. Measurements of cytokine secretion were made 8 and 24 hrs after activation in the presence of BSA (0.4 μM) using Max deluxe (Biolegend, CA, USA) ELISA kits [24]. Microglia were stimulated by bzATP (400 μM), LPS (1 μg/ml) and their combination. The protein content of the cells was measured using BCA Protein Assay (Pierce). Assays were repeated with internal control for cytokines for 5×10^5^-1×10^6^ cells per well. Cells were harvested by scraping with a rubber policeman, washed with PBS (4°C) and counted. We obtained ∼40 μg lysate from 5×10^5^ cells following lysis, clearance by centrifugation (14,000g, 10 min, 4°C). Each cytokine assay was calibrated by an internal standard curve.

### RNA-seq

Microglial cultures were harvested using a cell-scraper. Purification of total RNA was done for ∼10^6^ cells using QIAzol Lysis Reagent RNeasy plus Universal Mini Kit (QIAGEN, GmbH, Hilden, Germany). To ensure homogenization, a QIAshredder (QIAGEN, GmbH, Hilden, Germany) mini-spin column was used. Samples were transferred to a RNeasy Mini spin column and centrifuged for 15s at ≥8000g at room temperature. The mixture was processed according to the manufacturer’s standard protocol. Samples with an RNA Integrity Number (RIN) >8.5, as measured by Agilent 2100 Bioanalyzer, were considered for further analysis. Total RNA samples (1 µg RNA) were enriched for mRNAs by pull-down of poly(A)^+^ RNA. RNA-seq libraries were prepared using the KAPA stranded RNA-seq kit (Roche) according to the manufacturer’s protocol and sequenced using Illumina NextSeq 500 to generate 85 bp single-end reads.

### Bioinformatic analysis and Statistics

All next-generation sequencing data underwent quality control using FastQC [25] and were processed using Trimmomatic [26], aligned to GRCh38 using STAR. All genomic loci were annotated using GENCODE version 32 [27]. All experiments contained a minimum of three biological replicates. Trimmed Mean of M-values (TMM) normalization of RNA read counts and differential expression analysis were performed using edgeR [28]. All statistical tests were performed using R-base functions. Pathway and gene-set enrichment analysis were performed using the clusterProfiler: enrichKEGG program and the pathview for visualization. ID conversion from Ensemble to Entrez was carried out by the annotation package (biomaRt, org.Mm.eg.db). The partition of differential expressed (DE) genes to clusters was done according to threshold listed in **Table 1**.

**Table 1.**
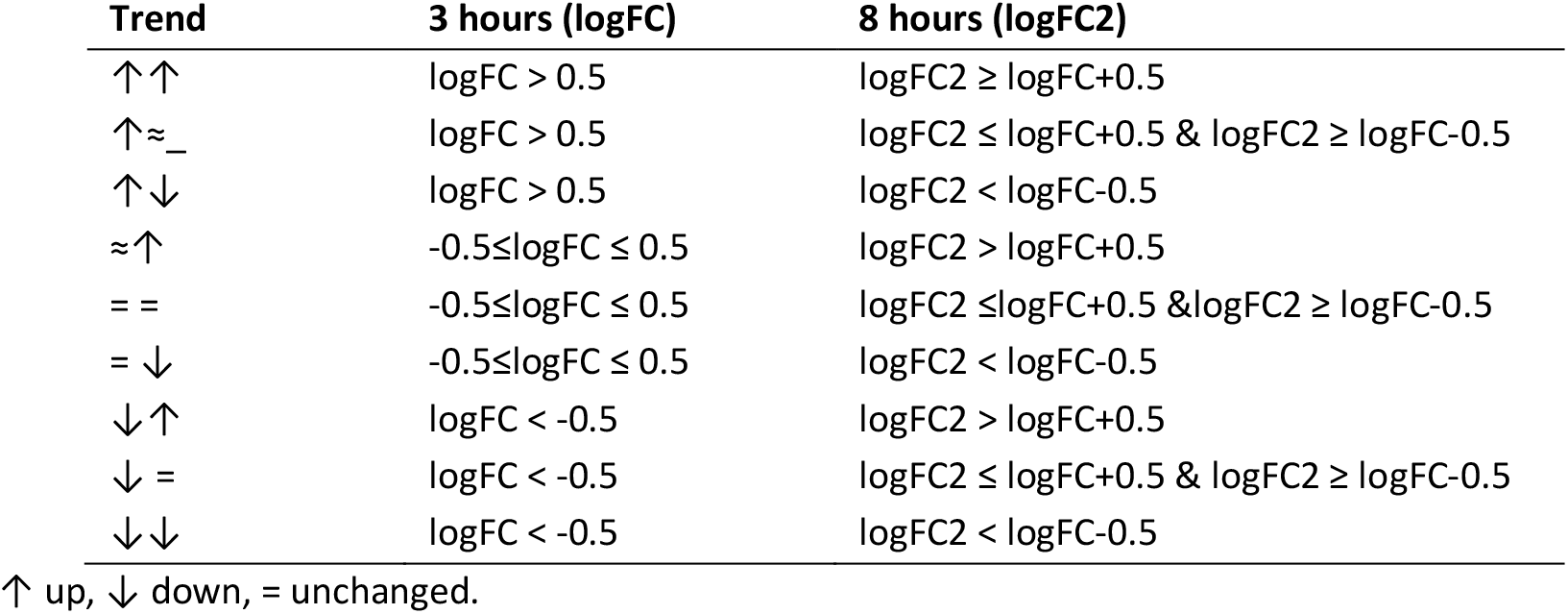
Thresholds used for assigning DE genes for 9 clusters with distinct expression trends

Experiments contained a minimum of three biological replicates. The cytokine quantification data were analyzed by one-way analysis of variance (ANOVA), using IBM SPSS Statistics Version 19 followed by Duncan’s *post hoc* test. Results from microglial cell experiments are presented as mean ± SD (standard deviation). When appropriate, p values <0.05 were calculated and considered statistically significant. Principal component analysis (PCA) was performed using the R-base function “prcomp”. Figures were generated using the ggplot2 R package.

## Results

### Functional response of primary microglia

The function of the microglia cells was monitored by quantifying the secretion of TNF-α and IL-6 following activation with bzATP, an agonist of purinergic P2X7 purinergic receptors plus LPS (**Figure 1**).

**Figure 1.**
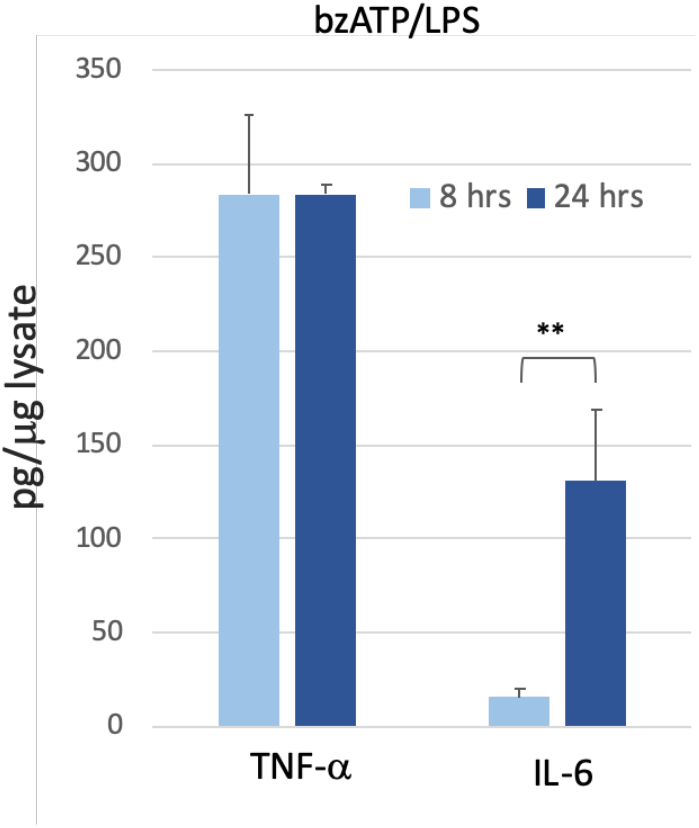
Quantitation of TNF-α and IL-6 released from primary neonatal murine microglial culture. Values are normalized with respect to the amounts of proteins (μg lysate) in the adhered cultured cells. The levels of TNF-α and IL-6 were below detection in naïve unstimulated cells. Samples for the analysis were derived from the conditioned media supplemented with BSA, harvested, and measured at 8 and 24 hrs after stimulation. Statistical significance marked by asterisks implies a p-value <0.02.

Secreted cytokines were below the level of detection prior to the addition of bzATP/LPS (<5 pg/μg of cell lysate) but increased markedly 8 and 24 hrs after simulation. While the amounts of IL-6 secreted were larger at 24 hrs than at 8 hrs (p value <0.02), those of TNF-α were stable, implying that the kinetics differ for each cytokine.

### A short-lived transient wave of gene expression by bzATP

To establish the characteristics of microglia and to quantify the molecular events that occur on changing cells from their resting state to maximal activation, we first tested the cells’ transcriptome at 3 hrs and 8 hrs following exposure to bzATP. Using RNA-seq analysis, we report 10,835 expressed genes that complied with the statistical quality threshold (see Materials and Methods). **Figure 2** shows the partition of these genes into 9 clusters along with their fold change (FC). The expression levels of almost all these genes (96%) is unchanged and thus are labeled ‘Same’ (**Figure 2A**). The term ‘Same’ is applied to changes in expression that are bounded by 50% (i.e., at a 0.67 to 1.5-fold difference relative to genes expressed in naïve cells). **Figure 2B** shows the partition of genes in the presence of bzATP for 3 hrs. Notably, at 8 hrs, the majority of the genes still marked as ‘Same’ (**Figure 2B**, gray color). The 9 expression groups are shown in **Supplemental Figure S1**.

**Figure 2.**
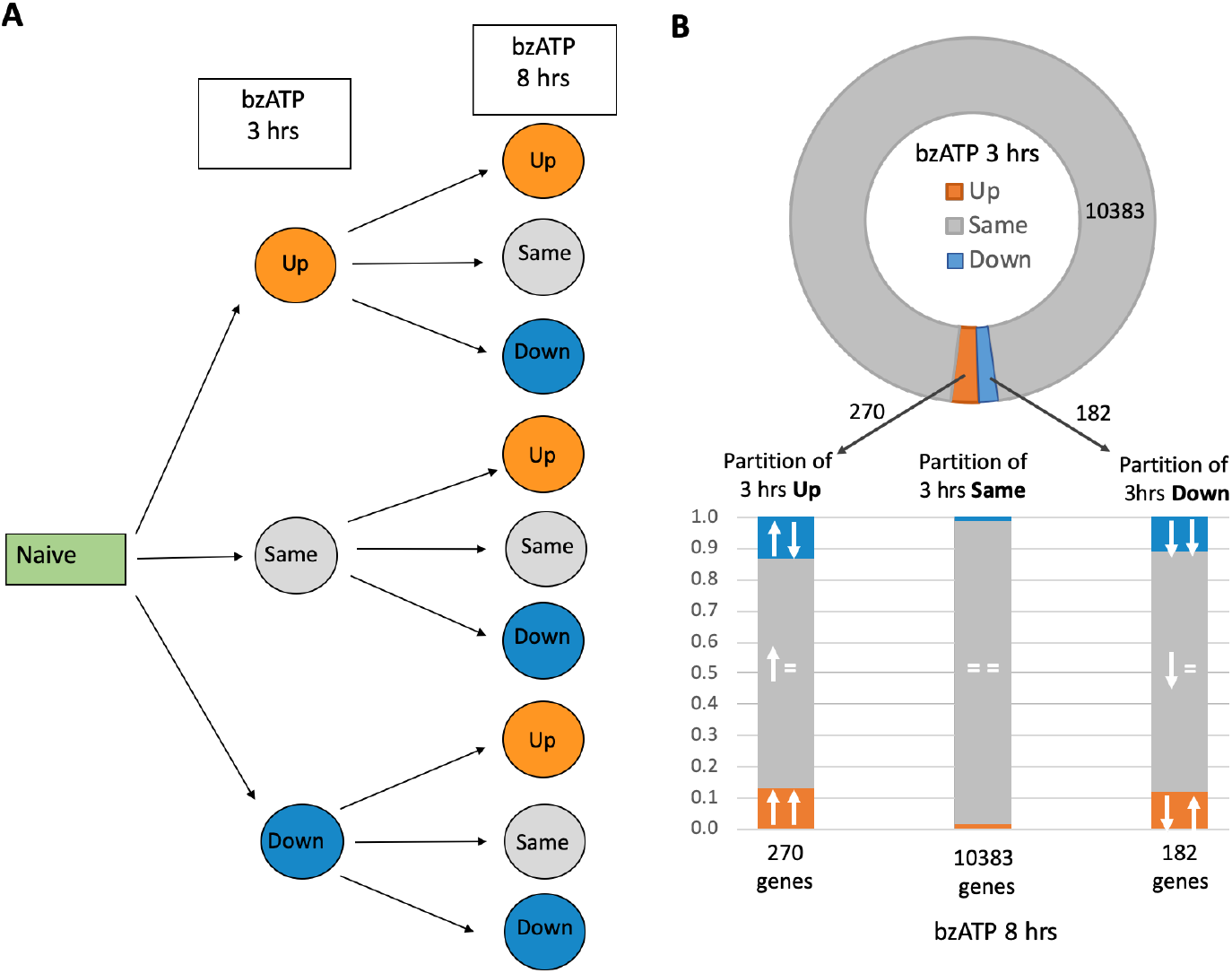
Trends in gene expression of microglial genes in the naïve (not treated) and following activation protocol with BzATP for 3 hrs and 8 hrs. **(A)** Differentially expressed genes are partitioned to 9 groups according to the joined expression trend (Up, Down or Same, and their combinations). **(B)** The ring-shaped graph shows the partition of the 10,835 expressed genes by their expression trend at 3 hrs of bzATP treatment relative to naïve cells. The number of genes labeled as Up, Same, Down is indicated. The histogram shows the partition of each of the three clusters according to the partition of expression trend measured at 8 hrs following bzATP exposure. Symbols for up (↑), down (↓) and same (=) are shown. The analysis is based on data in **Supplemental Table S1**.

Next, we tested the strength of the above observations by setting strict thresholds on the false discovery rate (FDR q-value < 0.05) and expression level (total >10 TMM for the average expression in naïve vs. bzATP treated cells), resulting in 33% of the original gene list (total of 3578 genes). Nevertheless, the vast majority of these genes (93%) are labeled ‘Same’, with only 16 genes (0.5%) significantly changing their expression in a monotonic trend (labeled Up-Up and Down-Down; **Supplemental Table S1**). Additionally, only 22 genes switched their trend of expression compared to that at 3 hrs. **Figure 3** shows the 7 genes that are downregulated (FDR q-value ranges from 3.1e-18 to 2.9e-10, **Figure 3A**), and 9 genes that are significantly upregulated (FDR q-value ranges from 2.7e-18 to 5.6e-04, **Figure 3B**). Among the strict Down-Down gene set, a cross-talk with the extracellular matrix is evident. For example, the enzyme hyaluronidase (Hyal1) and semaphorin receptor (Sema4d, also called Cd100) are involved in cell migration and differentiation.

**Figure 3.**
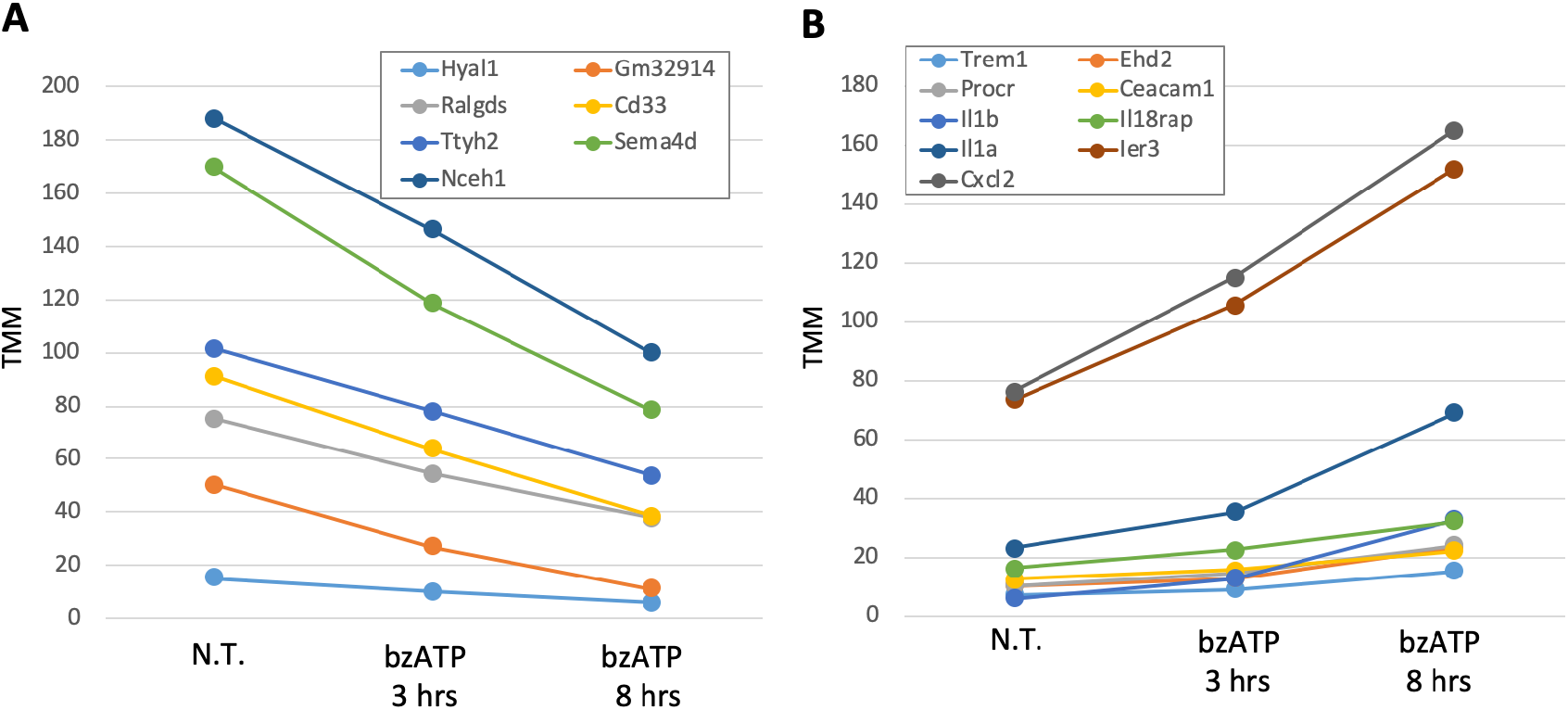
Dynamics of differential expressed (DE) genes following activation with bzATP. DE genes in naïve cells (N.T., not treated) and following bzATP treatment for 3 hrs and 8 hrs. The reported genes meet strict thresholds of FDR (q-value < 0.05) and expression level (total >10 TMM as an average expression of the naïve / bzATP treated cells). **(A)** Set of genes with a continuously reduced expression (labeled Down-Down). **(B)** Set of genes with upregulated expressed (labeled Up-Up). The analysis is based on data in **Supplemental Table S1**.

An inflammatory signal dominates the set of genes labeled Up-Up (9 genes, **Figure 3B**). Examples include Trem1, a scaffold membrane protein that acts in chemotaxis and regulates the killing of Gram-negative bacteria and Ceacam1, that is involved in signal transduction for T cell activation. The remaining genes belong to chemokine (Cxcl2, chemokine (C-X-C motif) ligand 2 and their immediate pathways (Il18rap, Il1a, IL1b). We conclude that exposing murine microglia to bzATP leads to changes in the levels of expression which are short-lived. This is shown by the transient wave of gene expression which fades during a longer time (8 hrs). Moreover, exposure to bzATP activates a distinctive set of genes that act as sensors and regulators of the immune axis of microglia.

### Genes that respond to bzATP with slow kinetics regulate microglial communication

In addition to DE genes that exhibit monotonic trends of expression (**Figure 3**), we investigated a set of genes with a slower change of kinetics (i.e., did not change significantly in 3 hrs but only after 8 hrs). **Table 2** lists the most significant of such DE genes (at a strict FDR <1.0e-10 for the kinetics of 3 hrs vs. 8 hrs; mean TMM>10, total 26 genes). It shows that many of the genes belong to trafficking and immune maintenance.

**Table 2.**
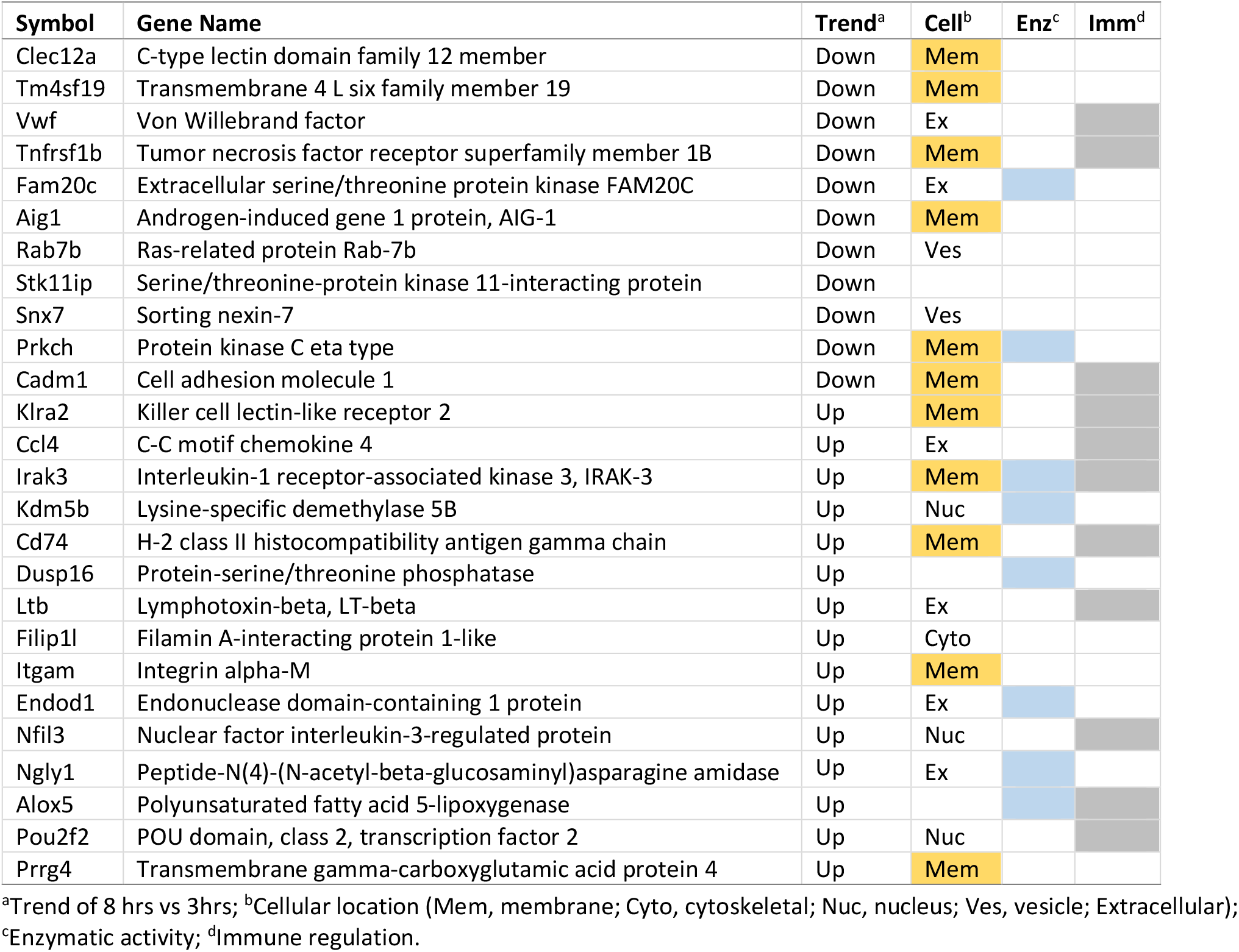
Differentially expressed genes by bzATP with slow kinetics

Among the DE genes marked as Same-Down are trafficking (i.e., Rab7b, Snx7) and cell interaction genes (e.g., Cadm1, Vwf). Cadm1 mediates homophilic cell-cell adhesion along with neuronal migration. Genes that were significantly changed with slow kinetics (**Table 2**) participate is the regulation of immune maintenance via TNF signaling. Tnfrsf1b (tumor necrosis factor receptor superfamily, 1b) and Ltb (Lymphotoxin-beta) promote cytokine cell surface signaling while Itgam directly activates TNF-primed neutrophils, regulates neutrophil migration and is involved in the production of superoxide ions in microglia. We conclude that the system is primed, but despite the strict parameters used for the late kinetic DE genes, the change in their expression is <2 fold (**Supplementary Table S2**).

### Expression of IL1 as an indicator of microglial activation

We characterized the function of cultured neonatal microglia by monitoring the release of three pro-inflammatory cytokines **(Figure 1)**. The set of interleukin (IL) genes and their receptors is a unique indicator for the state of brain inflammation. We assessed the relative expression of the transcripts of the IL-gene set (total of 33 expressed genes in microglia, collectively called IL-gene set) (**Supplementary Table S3**). The results from the RNA-seq from cells showed that IL-1β mRNA increases 4.2 fold within 3 hrs and 10.8 fold after 8 hrs exposure to bzATP (**Figure 4A**). Despite this high increase in expression, IL-1β accounts for only 2% of the total TMM of the IL gene set (considered expressed IL-gene set as 100%).

**Figure 4.**
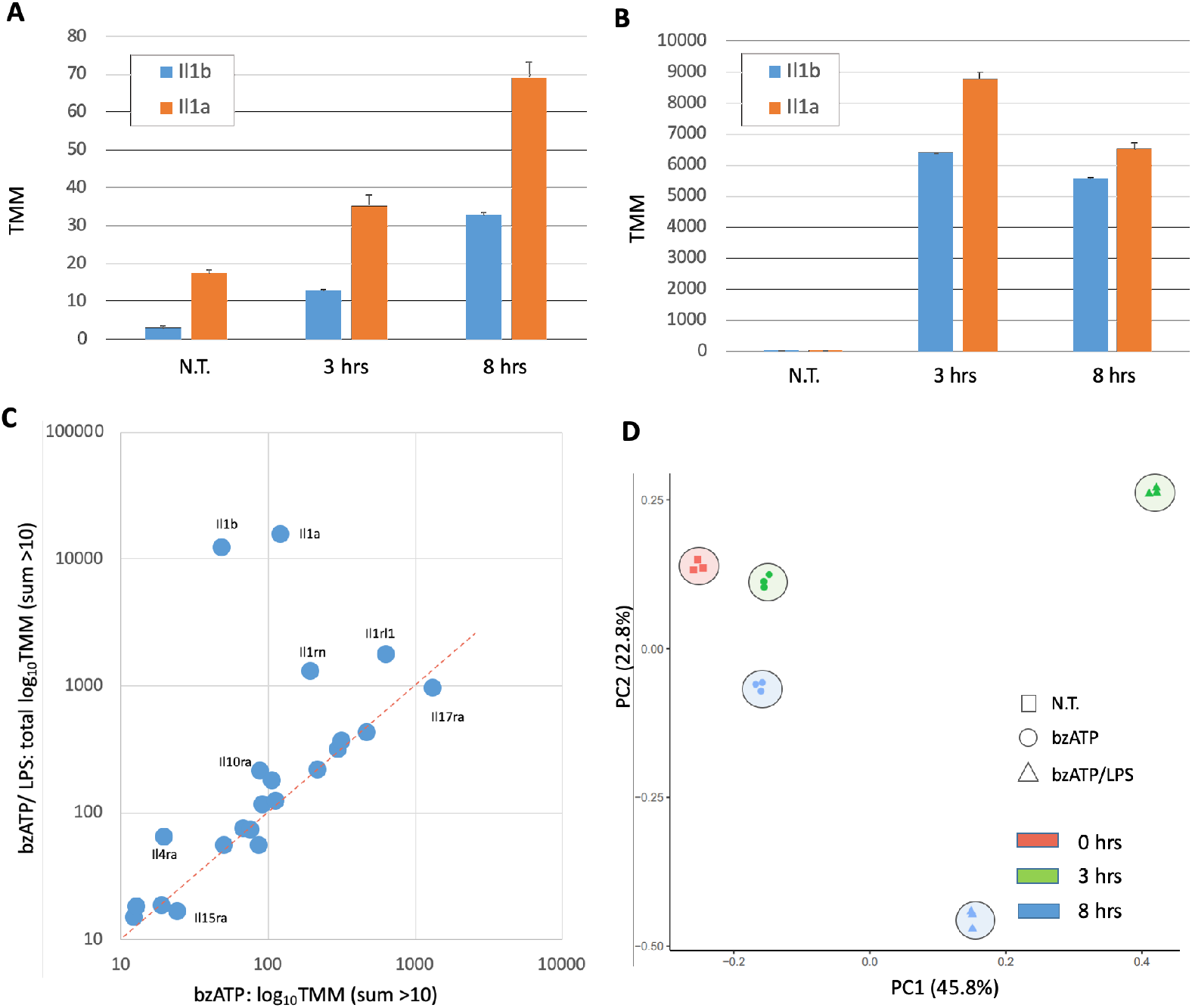
Dynamics of differential gene expression of interleukins and their direct regulators (IL-gene set) in naïve (N.T., not treated) and following treatments for 3 hrs and 8 hrs. **(A)** The expression levels of Il1a and Il1b following activation with bzATP. **(B)** The expression levels of Illa and Il1b following activation with bzATP/LPS. Note the difference in the y-axis. The SDs (standard deviations) are based on the triplicates from RNA-seq. **(C)** Analysis of 33 IL-gene set under activation condition by bzATP (x-axis) and bzATP/LPS following 8 hrs (y-axis). Only genes with mean expression levels >10 TMM are shown. **(D)** PCA analysis of the 15 samples of RNA-seq. The variance explained by the two principle components (PC1, PC2) is 68.6%.

By contrast, after the addition of LPS to bzATP, IL-1β mRNA levels increase ∼2100 fold within 3 hrs (**Figure 4B**) and account for 35% of the expressed mRNAs of interleukins and their receptors (i.e., IL-gene set) and 39%, at 8 hrs. The kinetics of Il1a is similar to that of Il1b. bzATP plus LPS increases IL-1α mRNA levels ∼500 fold within 3 hrs and the IL-1α transcript accounts for 48% of the total interleukin mRNAs in such activated cells. Unlike interleukin itself, the receptor Il1rl1 explains 10% of the interleukin expressed gene set in untreated cells and increases 9 fold, 3 hrs after activation by BzATP/LPS with no further increase in RNA level at 8 hrs. (**Figure 4C**).

We found that IL-1 regulators are also strongly responsive to the culture conditions. The expression of the Interleukin 1 receptor antagonist (Il1rn), which inhibits the activities of IL-1α and IL1β is transiently induced by bzATP (1.8 folds, 3 hrs), but returns to its baseline levels at 8 hrs. The addition of LPS leads to a substantial and permanent increase in its expression (13.1 and 10.7 folds in 3 and 8 hrs, respectively, **Figure 4C**).

LPS also changed the gene expression of IL-4 and IL-10 receptors by >4 fold (**Figure 4C; Supplementary Table S3**). In the naïve cells, Il17ra accounts for 30% of interleukin transcripts. They are increased 25% by exposure to bzATP and reduced by the addition of LPS (25% of its basal level; **Figure 4C**). Il17ra (interleukin 17 receptor A) showed an inverse trend relative to the discussed IL genes. Based on the expression trend of the IL-17 receptor, we postulate that it is not part of the cell activation program in primary microglial cells.

We tested the transcriptional programs from RNA-seq of all 15 tested samples (5 experimental groups, see Materials and Methods) by applying PCA analysis (**Figure 4D**). Triplicates (sharing the same color and symbol) are clustered tightly confirming that the variance within a group is negligible relative to the variance of the tested conditions (i.e., NT, bzATP, and bzATP/LPS in two time points). While the actual position in the 2-dimensional representation is somewhat arbitrary, the PCA is consistent with an effect for bzATP that is relatively restricted, in contrast to the strong and persistent effect induced by bzATP/LPS. Moreover, the delayed kinetics allows consolidation of the changes of the entire transcriptome. The variance explained by the two principle components (PC1, PC2) is 68.6%.

### Global alteration of cell transcriptome by bzATP/LPS is synchronized and long-lasting

**Figure 5A** shows the partition of the DE genes (total of 10,769 genes) into 9 clusters in the presence of bzATP/LPS. The expression trends for all 10,769 genes is partitioned to 3 cluster, labelled Up, Same and Down. A marked difference is seen from activation by bzATP alone (compare **Figure 2A** to **Figure 5A**). **Figure 5A** shows that the a large fraction of the genes (44.3%) is drastically changed in expression by bzATP/LPS within 3 hrs (1924, 2851 for Up and Down labeled genes, respectively). The dynamics of all 10,769 genes before, (time=0) and 3 hrs and 8 hrs after activation is shown (**Supplemental Figure S4**).

**Figure 5.**
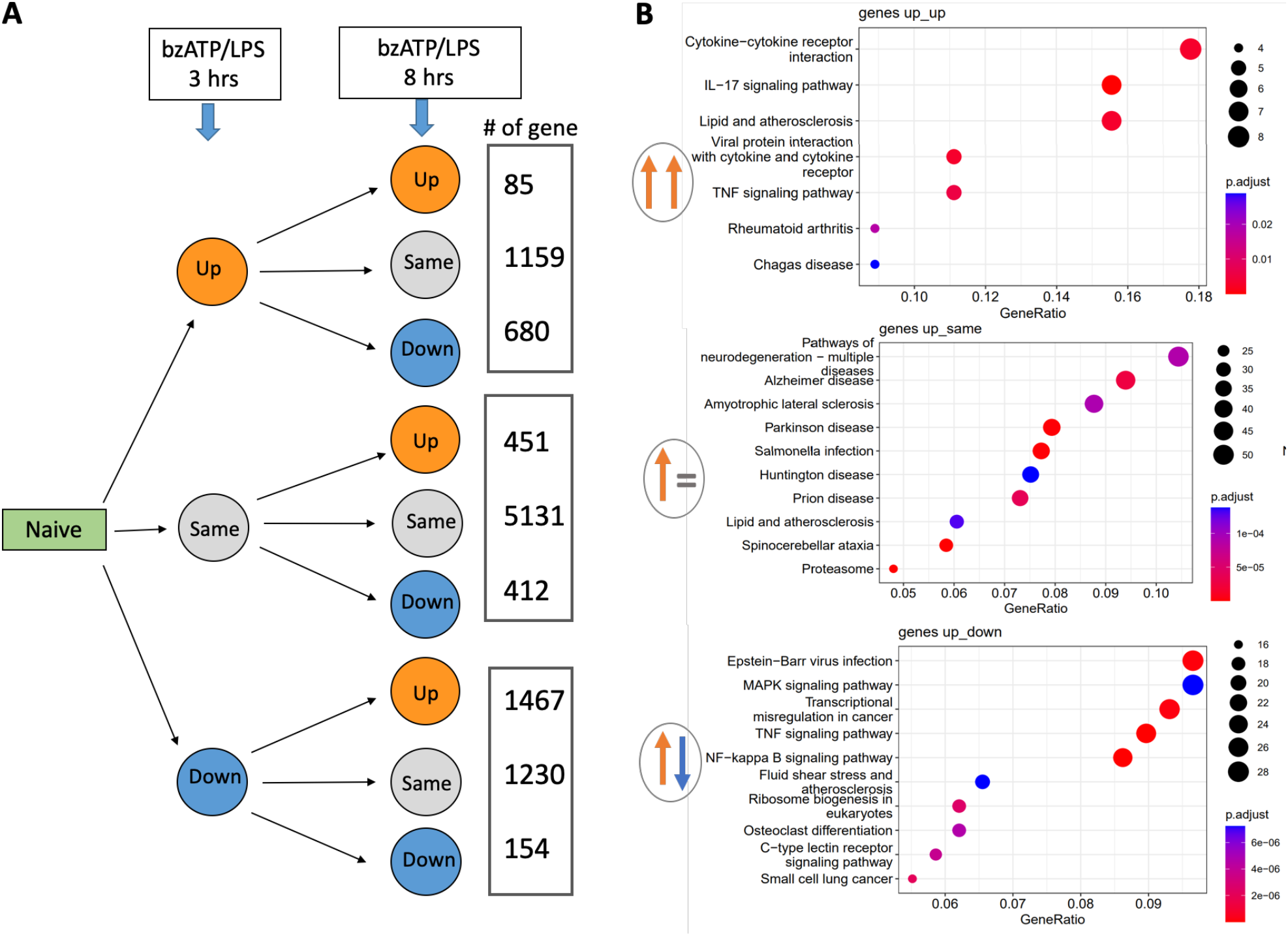
Partition of genes by their expression trends and their functional enrichment. **(A)** Gene expression levels of naïve microglial cells and following activation with bzATP/LPS for 3 hrs and 8 hrs. A scheme of all 9 expression trends for 10,769 genes together with the number of genes associated with each cluster is shown. For details see **Supplementary Table S4. (B)** Enrichment analysis for DE genes for genes induced after 3 hrs (i.e., labeled Up). Annotations are according to the KEGG pathway resource. Statistical enrichment of p-adjust is depicted by the colors (blue to red) for FDR <0.05. The size of the dots captures the number of proteins. Note that the number of genes associated with each enriched pathway, and the gene ratio (i.e., the fraction of genes in the cluster that are included in a specific pathway, x-axis) differ among gene clusters. The pathway enrichment analysis for all 9 expression trend clusters is shown in **Supplemental Figure S3**.

**Figure 5B** shows the enrichment of cellular pathways (by KEGG) for the genes that were induced by bzATP/LPS at 3 hrs. The partition of these genes (total 1924) by their dynamic expression trend (Up-UP, Up-Same, and Up-Down) is shown in **Figure 5B**. In the cluster Up-Up (85 genes), the listed pathways are associated with cytokines (e.g., TNF signaling), viral infection, and inflammation-based diseases. By contrast, Up-Same (1159 genes) cluster shows enrichment in aging and degenerative CNS diseases, and the Up-Down (680 genes) cluster is enriched in signaling of MAP kinase, TNF, and NFκB. Results of an enrichment analysis for all DE genes to known cellular pathways for all 9 clusters are shown in **Supplemental Figure S2**. We conclude that the strong signal on immunological response, neurodegenerative diseases, and immune-response signaling pathways substantiate this culture as a responsive cellular system with a robust transcriptional reactivity.

We then tested genes that were upregulated by bzATP/LPS through their gene ontology (GO) annotations (**Figure 6)**. We focused on using 585 statistically significant DE genes that were strongly induced (>1.5-fold increase in expression relative to that of non-treated cells and an average expression level >10 TMM). Inspection of enrichment in biological processes (**Figure 6**, left) and molecular function (**Figure 6**, right) revealed that the significant enrichment of the GO annotations and their interconnection is dominated by an immunological signature. The most significant biological processes listed are cytokine and chemokine regulation and immunological response. The results from analysis with a more relaxed threshold (p-value <1.0e-07) for individual GO annotation are shown in **Supplementary Figure S4**. It includes a signal for phosphorylation regulation for GO molecular function. We conclude that an unbiased view of gene expression supports a robust and long-lasting induction of signaling of cytokines and chemokines.

**Figure 6.**
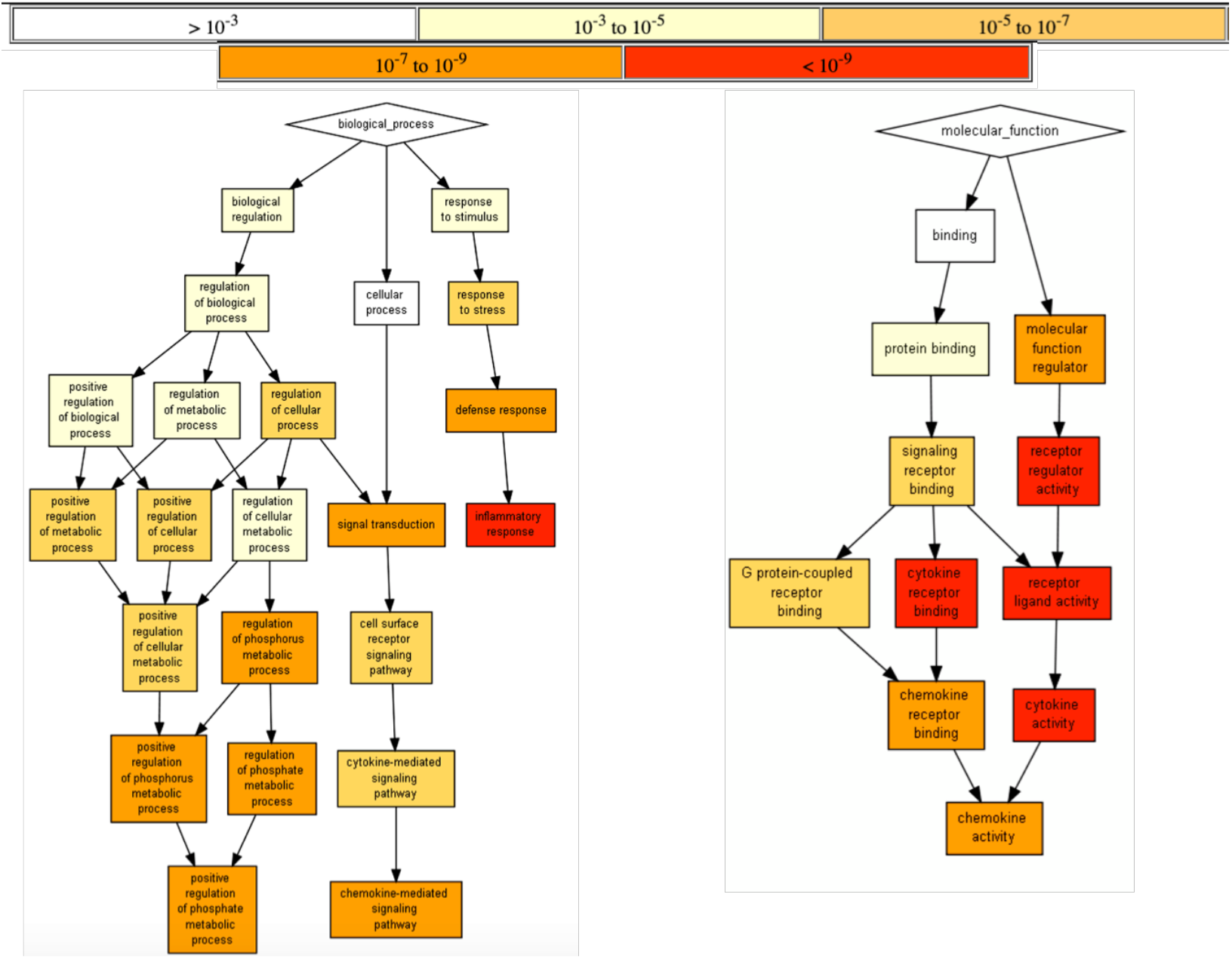
Enrichment analysis for gene ontology (GO) of upregulated DE genes. An enrichment test was applied on 585 statistically significant DE genes that were strongly induced (>1.5-fold increase in expression relative to that of non-treated cells, and average expression level >10 TMM). The test was based on hypergeometric analysis, followed by multiple testing corrections, (GOrilla statistical tool [29]). Results shown are for biological process (left) and molecular function (right). Only GO terms with p-value <1.0e-08 were included in the analysis. Top, statistical significance color-coded range for the calculated adjusted-p-values for the GO annotations. The color scale (white to red) with the associated range of p-values is shown (Top).

### Changes in gene expression by orders of magnitude drive microglia activation state

**Figure 7** shows a sorted list of 25 genes that were selected by the scale of induction relative to naïve cells for 8 hrs in the presence of bzATP/LPS. Despite the transient signal following bzATP, the scale of the induction is already very large at 3 hrs and remains for most genes also at 8 hrs (**Figures 7A-7B**). The induction ranges across 5.5-2800 fold **(Figure 7C)**. The majority of the induced gene products are functionally and physically connected **(Figure 7D)**. Among the highly connected proteins are Tnf and Il1b followed by components of the chemokine signaling (Cxcl10, Ccl2, Cxcl2, Ccl5, Csf2). Some of the induced chemokines are known in the context of neuronal survival. For example, Ccl5 activates downstream signaling pathways involving MAP kinases.

**Figure 7.**
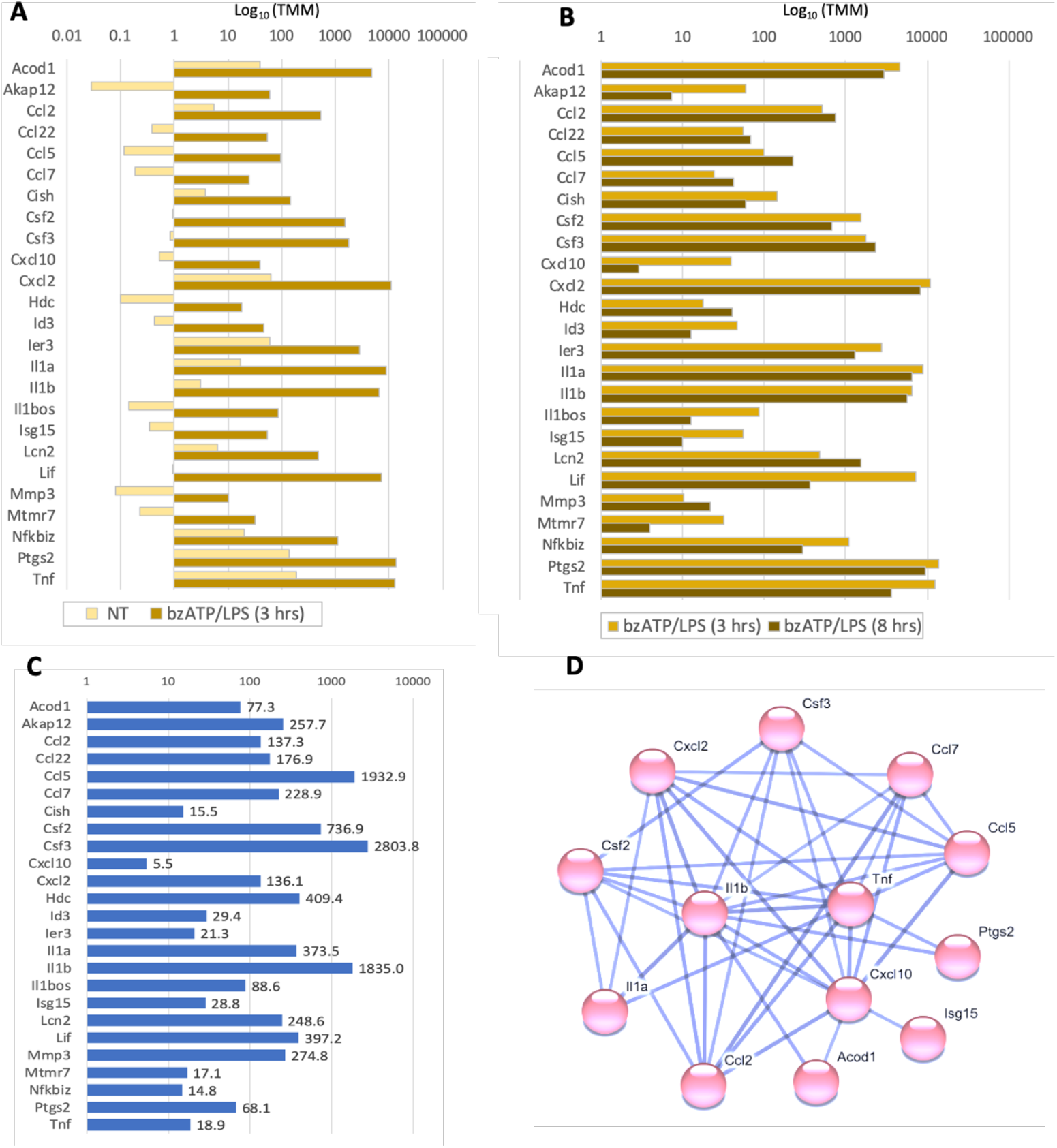
Top 25 induced genes sorted by their scale of induction by bzATP/LPS after 8 hrs relative to expression in naïve cells. **(A)** log_10_TMM of the expression in naïve, non-treated cells (NT) and 3 hrs of bzATP/LPS. **(B)** log_10_TMM of the expression in 3 hrs of bzATP/LPS and 8 hrs of bzATP/LPS. The genes in A-C are listed alphabetically. **(C)** Increase at 8 hrs relative to that in naïve cells, e.g., Csf3 (colony stimulating factor 3) increases 2803.8 fold. For details see **Supplementary Table S5. (D)** A connectivity map according to STRING [30]) protein interaction network. Only the genes (13/25) with high confidence connection score >0.9, are shown.

Applying an unbiased enrichment analysis for the most significant downregulated genes (with >10 TMM and suppressed expression by a factor of 2-20, total 720 genes). Inspecting these genes revealed no significant enrichment in the GO annotation analysis, functional connectivity or KEGG pathways (**Supplementary Figure S3**). Of a special interest is P2rx7, the ATP-gated cation channel that participates in neuroinflammation and pathophysiological processes. P2rx7 is already downregulated 8.3 fold at 3 hrs after the addition of bzATP/LTP. It was shown that P2rx7 mRNA is suppressed when cells are exposed to inflammatory conditions and microglial cell death is induced by bzATP [24]. Between 3 to 8 hrs the P2rx7 transcript is induced 3 fold, following the transient suppression at the earlier time point. We conclude that in stimulated microglial cells, hundreds of genes are upregulated and downregulated but only those that are induced by several orders of magnitude are coordinated to execute the robust inflammatory signature.

### Alternative routes for activation state by different cell surface signaling patterns

We compared our results with RNA-seq from that in isolated microglia from C57BL/6J mice before and after activation for 6 hrs with LPS+IFNγ [31]. Several of the highest upregulated genes (e.g., Tnf, Csf3, **Figure 7**) showed a similar trend of induction in the two experiments. To pinpoint the subtle difference in signaling in both settings, we focused on TNF signaling pathways (KEGG [32]) and reanalyzed the DE genes. **Figure 8** shows the expression trend following 8 hrs in bzATP/LPS compared to activation by LPS+IFNγ [31]. The TNF signaling pathway shows a signaling cascade that starts with the binding of TNFα through the execution of cell survival, apoptosis, necroptosis, the secretion of interferon β, and the transcription of immune genes including inflammatory cytokines. We highlight strong differences in expression trend along the signaling cascade (**Figure 8**, yellow arrows).

**Figure 8.**
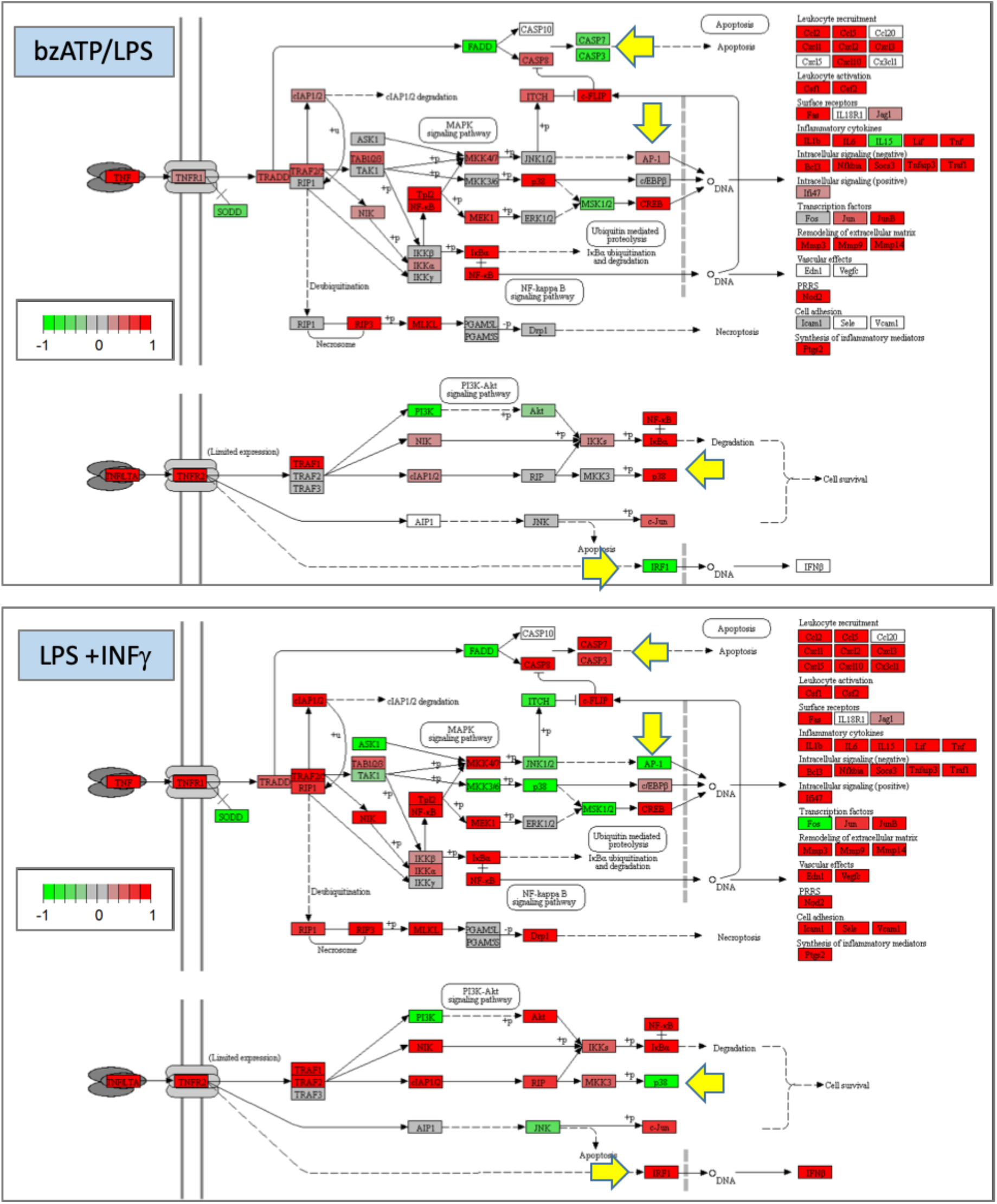
KEGG pathway map of TNF signaling. The top panel indicates the genes that are downregulated (green) upregulated (red), unchanged (Gray), and undetected (white). The DE genes in cells exposed to bzATP/LPS (8 hrs, Top), and LPS+IFNγ for 6 hrs (Bottom) are shown. Selected genes for which expression trends between the two settings of microglia activation substantially differ are marked with yellow arrows.

While apoptosis is induced by activation with LPS+IFNγ (see Casp3, Casp7), these genes are suppressed by bzATP/LPS. Moreover, bzATP/LPS does not change the expression of RIP1 but it is upregulated in microglia exposed to LPS+IFNγ. RIP1 is a kinase that acts to balance cell death and survival [33]. It is likely that for activation by bzATP/LPS, the signaling cascade shifts towards NF-κB branch that initiates MAP kinase signaling. Activation with bzATP/LPS causes overexpression of p38. All other components in this signaling are either unchanged or upregulated. The expression pattern of the MAP kinase genes under the condition of LPS+IFNγ indicates that MKK3/6, p38, JNK1/2, and ITCH genes are downregulated which eventually leads to the downregulation of AP-1 (activator protein 1), whose level of expression is fundamental to the induction of the innate immune response. Many gene products are listed as the outcome of TNF signaling (**Figure 8**, right). Notable, the expression of several of these genes is undetected in cells exposed to bzATP/LPS.

Among these are leukocyte recruitment (Ccc15, Cx3cl1), cell adhesion (Sele, Vcam1) and vascular effect (Edn1, Vegfc) gene sets. In addition, the inflammatory cytokine IL15 is downregulated only by bzATP/LPS. Suppression of IL15 in a microglial culture induced by LPS+IFNγ led to a defective proinflammatory response, suggesting its pivotal role in the feedback and signaling of the immune response [34]. Of the transcription factors, Irf1 is a likely a regulator of various pro- and anti-inflammatory states [35]that regulates the release of IFN-β. Only in microglia activated by LPS+IFNγ but not by bzATP/LPS, is the expression of both Irf1 and INF-β coordinately upregulated. Irf1 rapidly enhanced IFN-β and IFN-λ after stimulation with nucleic acids that mimic viral infection (poly I:C), thereby inducing IFN stimulated genes [36, 37]. We conclude that the robust programs of activation of microglia with different combinations of stimuli resulted in a rich repertoire of distinct responses that define the underlying biology of the inflammatory routes induced in microglial cells.

## Discussion

Microglia switch between resting and activated stages to fulfill their surveillance and homeostatic function in the brain [38]. They are characterized by their capacity for sensing abnormal and disturbed conditions that may originate from toxic agents, physical injury, pathogens and in the course of brain aging [39]. Because of the crucial importance of microglia in maintaining brain homeostasis, the kinetics and amplitude of their response must be tightly controlled. In this study, we searched for a molecular characterization underlying the response of a microglial neonatal culture to two external stimulants.

The induction of microglia starts with the binding of extracellular molecules to Toll-like receptors [40]. The outcome *in vivo* and *in vitro* results in changes in cytokine secretion, cell proliferation, migration, and phagocytosis [41]. *In vivo*, microglia respond by sensing the local concentrations of molecules, including amyloid β (Aβ), nucleic acids, reactive oxygen species (ROS) and ATP. The latter is released from dying cells after a stroke or injury [42, 43]. Release of ATP from astrocytes and neurons to the extracellular space triggers microglia to change their activity [44, 45]. Microglia also release ATP through exocytosis [4], thereby joining a feedback loop to modulate the activation program. The role of bzATP for priming microglial cells is of especial importance in an *in vitro* model for the study of neuropathology [46].

In the current study, we used a purified primary microglial culture to examine the effect of extracellular ATP (in the form of BzATP) that acts on purinergic P2rx7 receptors on the cell membrane [47]. By maintaining microglia under controlled conditions, we found that the variance in gene expression levels is negligible in samples from the same experimental conditions (see PC analysis; **Figure 4C**), confirming the high confidence of studying the expression trend of specific genes. We show that exposing cells to bzATP produces an immediate and synchronized response (**Figures 2-3**). However, only following a combination of bzATP with LPS, is the response consolidated and becomes a long-lasting stronger signal affecting hundreds of genes (**Figures 5-8**). Presumably, priming by bzATP drives microglia to become more sensitive to other stimuli (e.g., LPS). However, if such a stimulus is delayed or fails to reach an activation threshold, the transcriptional wave fades. We observed that 3.1% of the genes (from 10,835 genes) changed significantly 3 hrs after exposing cells to bzATP, but most of them returned to their baseline levels after 8 hrs. Genes that are strongly induced by bzATP at 3 hrs include Traf5 (1.6 fold) and Malt1 (1.4 fold), proteins known to affect NF-κB signaling through a ubiquitination regulation [48].

A strong 2.2 fold upregulation of Ccl2 (chemokine C-C motif ligand 2), (p-value 9.3e-14) is of a special interest due to its relevance to brain pathology and it is upregulated in the brain in AD disease [49]. In mouse models of AD, the overexpression of Ccl2 was shown to cause microglia-induced amyloid β oligomerization, worsening of tau pathology and an increase in IL-6 [50]. The transient wave of bzATP-driven DE genes highlights the relevance of the innate immune system and the role of microglia in brain pathogenesis. It can provide a baseline for testing drugs in varying conditions [51].

The molecular signature of cell activation has been extensively studied in the BV2 microglial cell line [52]. However, the transcriptomes of BV2 and the primary microglial cells [53, 54] are quite different. We show that the outcome of activated primary microglial culture is a steady secretion of cytokines (**Figure 1**). The induction of cytokine and chemokine DE genes in the presence of bzATP/LPS is coordinated in time and reach an increase in amounts by several orders of magnitude **(Figures 4, 7**). We show that even in an isolated culture, the responsiveness of the cells remains high despite the lack of neurons, astrocytes, or other cell types in the sorounding, indicating the intrinsic potency of the primary microglia culture.

We aimed to distinguish two alternative models for microglial activation: (i) Activated cells are programed to converge into a defined transcriptomic profile irrespective of the nature of the stimulus; (ii) The cells can be stabilized in distinct inflammatory states, each of which displays a unique transcriptional profile according to the induced physiological scenario. Accordingly, the concentration, duration and identity of the stimulus (e.g., bzATP, IFNγ, TNF, etc.) define the outcome. Our results support the second model in which a large variability exists among experimental conditions but not within any tested condition **(Figure 4D**).

The RNA-seq data for activation of neonatal microglia mimic that of cells exposed to dying neuronal cells (bzATP), viral infection (IFNγ), and pathogens (LPS). Microglia exposed to LPS+IFNγ for 6 hrs [31] simulate the inflammatory response upon viral infection. IFNγ is a strong mediator for a wave of immune-related gene expression [55]. **Figures 8** compares the studies on the TNF signaling pathway. The analysis of the RNA-seq data captures the expression trend of each of the studies separately, together with that of TNF signal transduction. Differences in the concentration of LPS and time of measurements (8 hrs in bzATP/LPS and 6 hrs in LPS+IFNγ [31] did not substantially change the overall response. In both cellular settings, the overexpressed genes account for the majority of the signal (**Figures 8A-8B**, red). Nevertheless, the fraction of genes that are labeled ‘same’ (i.e., stable expression, unaffected by bzATP/LPS) varies between these studies, with a 3-fold higher number of such genes for bzATP/LPS than for the LPS+IFNγ condition.

We questioned whether this observation is specific to TNF signaling pathway and tested another inflammation-related signaling pathway of NF-κB (**Supplementary Figure S5**). NF-κB is a hub for executing the inflammatory signal in microglia and macrophages (e.g., RAW 264.7 cells) [56]. It is a master regulator linking LPS with the production of IL-10 and IL-6 [57]. We observed that while only 16.5% of the genes overlap between the TNF and NF-kB signaling pathways, the overall partition of genes according to their expression trend shows that 2.8-fold more genes are marked as unchanged for bzATP/LPS than for LPS+IFNγ (labeled Same, 28 vs. 10 genes) in the NF-kB signaling pathway.

In the aging brain, different stimuli that mimic injury, hypoxia, and pathogenic conditions are associated with inflammatory cytokines (e.g., TNF-α, IL-6, and IL-1β). In the postmortem brain tissue of individuals with AD, [58], PD, or Multiple Sclerosis) [59], neuroinflammation is considered as the main driver of the disease. Inflammation mediated by microglia, together with damaging oxidative stress accelerates in the progression of neurodegeneration [14]. Consequently, inhibiting microglial activation should be a strategy for slowing down the progression of the disease in the elderly population. We have recently shown that ladostigil, a drug with anti-oxidant and anti-inflammatory activity has a beneficial effect on the memory decline in the aged rat brain [20]. We found that the number of microglia and their morphology is significantly changed in a region-dependent manner in aging rats chronically treated with ladostigil [60]. We propose adopting the *in vitro* neonatal primary culture as a model system because it is sensitive to cues in the microenvironment. Such a system is attractive for unveiling the underlying mechanism of anti-inflammatory drugs and in the attempts to manage pathologies of the CNS in neurodegenerative diseases.

## Abbreviations

AD: Alzheimer’s disease
ATP: Adenosine tri-phosphate
BSA: Bovine serum albumin
CNS: Central nervous system
DE: Differentially expressed
DMEM: Dulbecco’s modified Eagle medium
FC: Fold change
FDR: False discovery rate
GO: Gene ontology
Hrs: Hours
IL: Interleukin
IFNγ: Interferon gamma
LPS: Lipopolysaccharide
NT: Not treated
PCA: Principal component analysis
PD: Parkinson’s disease
RNA-seq: RNA sequencing analysis
SD: Standard deviation
TMM: Trimmed mean of means

## Supplementary Tables

**Table S1**. RNA-seq results for bzATP treatment (by TMM)

**Table S2**. Gene set of DE genes exposed to bzATP with slow kinetics (Source of Table 2)

**Table S3**. RNA-seq results for 33 interleukin-related genes (IL-gene set; Source of Figure 4C).

**Table S4**. RNA-seq results for bzATP/LPS treatment (by TMM)

**Table S5**. Top 25 induced genes (Source of Figure 7).

## Supplemental Figures

**Figure S1**. Gene expression trends on naïve, not treated microglial cells and following activation with bzATP for 3 hrs and 8 hrs.

**Figure S2**. Gene expression trends on naïve, not treated microglial cells and following activation with bzATP/LPS for 3 hrs and 8 hrs.

**Figure S3:** Enrichment analysis for all 9 differentially expressed (DE) gene clusters followed bzATP/LTP induction.

**Figure S4**. Enrichment analysis for gene ontology (GO) of upregulated DE genes.

**Figure S5**. DE gene pattern on the KEGG pathway map of NF-κB signaling.

## Acknowledgements

We would like to thank the lab members for useful comments and fruitful discussion. This study was supported by

## Ethical

All animals related studies were performed according to the National Research Council’s guide and approval by the Hebrew University Institutional Committee.

## Data availability

RNA-seq data files were deposited in ArrayExpress under the accession <u>E-MTAB-10450</u>

## References

1. Amor, S.; Peferoen, L. A.; Vogel, D. Y.; Breur, M.; van der Valk, P.; Baker, D.; van Noort, J. M., Inflammation in neurodegenerative diseases–an update. Immunology 2014, 142, (2), 151–166.

2. Clayton, K. A.; Van Enoo, A. A.; Ikezu, T., Alzheimer’s disease: the role of microglia in brain homeostasis and proteopathy. Frontiers in neuroscience 2017, 11, 680.

3. Gao, H.-M.; Hong, J.-S., Why neurodegenerative diseases are progressive: uncontrolled inflammation drives disease progression. Trends in immunology 2008, 29, (8), 357–365.

4. Imura, Y.; Morizawa, Y.; Komatsu, R.; Shibata, K.; Shinozaki, Y.; Kasai, H.; Moriishi, K.; Moriyama, Y.; Koizumi, S., Microglia release ATP by exocytosis. Glia 2013, 61, (8), 1320–30.

5. Sanz, J. M.; Di Virgilio, F., Kinetics and mechanism of ATP-dependent IL-1 beta release from microglial cells. J Immunol 2000, 164, (9), 4893–8.

6. Neher, J. J.; Neniskyte, U.; Brown, G. C., Primary phagocytosis of neurons by inflamed microglia: potential roles in neurodegeneration. Front Pharmacol 2012, 3, 27.

7. Fricker, M.; Vilalta, A.; Tolkovsky, A. M.; Brown, G. C., Caspase inhibitors protect neurons by enabling selective necroptosis of inflamed microglia. J Biol Chem 2013, 288, (13), 9145–52.

8. Hammond, T. R.; Dufort, C.; Dissing-Olesen, L.; Giera, S.; Young, A.; Wysoker, A.; Walker, A. J.; Gergits, F.; Segel, M.; Nemesh, J., Single-cell RNA sequencing of microglia throughout the mouse lifespan and in the injured brain reveals complex cell-state changes. Immunity 2019, 50, (1), 253–271. e6.

9. Park, J.; Min, J.-S.; Kim, B.; Chae, U.-B.; Yun, J. W.; Choi, M.-S.; Kong, I.-K.; Chang, K.-T.; Lee, D.-S., Mitochondrial ROS govern the LPS-induced pro-inflammatory response in microglia cells by regulating MAPK and NF-κB pathways. Neuroscience letters 2015, 584, 191–196.

10. Cobourne-Duval, M. K.; Taka, E.; Mendonca, P.; Soliman, K. F., Thymoquinone increases the expression of neuroprotective proteins while decreasing the expression of pro-inflammatory cytokines and the gene expression NFκB pathway signaling targets in LPS/IFNγ-activated BV-2 microglia cells. Journal of neuroimmunology 2018, 320, 87–97.

11. Wang, W. Y.; Tan, M. S.; Yu, J. T.; Tan, L., Role of pro-inflammatory cytokines released from microglia in Alzheimer’s disease. Ann Transl Med 2015, 3, (10), 136.

12. Rodriguez-Gomez, J. A.; Kavanagh, E.; Engskog-Vlachos, P.; Engskog, M. K. R.; Herrera, A. J.; Espinosa-Oliva, A. M.; Joseph, B.; Hajji, N.; Venero, J. L.; Burguillos, M. A., Microglia: Agents of the CNS Pro-Inflammatory Response. Cells 2020, 9, (7).

13. Welser-Alves, J. V.; Milner, R., Microglia are the major source of TNF-alpha and TGF-beta1 in postnatal glial cultures; regulation by cytokines, lipopolysaccharide, and vitronectin. Neurochem Int 2013, 63, (1), 47–53.

14. Simpson, D. S. A.; Oliver, P. L., ROS Generation in Microglia: Understanding Oxidative Stress and Inflammation in Neurodegenerative Disease. Antioxidants (Basel) 2020, 9, (8).

15. Smith, J. A.; Das, A.; Ray, S. K.; Banik, N. L., Role of pro-inflammatory cytokines released from microglia in neurodegenerative diseases. Brain research bulletin 2012, 87, (1), 10–20.

16. Cianciulli, A.; Porro, C.; Calvello, R.; Trotta, T.; Lofrumento, D. D.; Panaro, M. A., Microglia Mediated Neuroinflammation: Focus on PI3K Modulation. Biomolecules 2020, 10, (1).

17. Zhou, L.-t.; Wang, K.-j.; Li, L.; Li, H.; Geng, M., Pinocembrin inhibits lipopolysaccharide-induced inflammatory mediators production in BV2 microglial cells through suppression of PI3K/Akt/NF-κB pathway. European journal of pharmacology 2015, 761, 211–216.

18. Noristani, H. N.; Gerber, Y. N.; Sabourin, J.-C.; Le Corre, M.; Lonjon, N.; Mestre-Frances, N.; Hirbec, H. E.; Perrin, F. E., RNA-seq analysis of microglia reveals time-dependent activation of specific genetic programs following spinal cord injury. Frontiers in molecular neuroscience 2017, 10, 90.

19. Hirbec, H.; Marmai, C.; Duroux Richard, I.; Roubert, C.; Esclangon, A.; Croze, S.; Lachuer, J.; Peyroutou, R.; Rassendren, F., T he microglial reaction signature revealed by RNA seq from individual mice. Glia 2018, 66, (5), 971–986.

20. Linial, M.; Stern, A.; Weinstock, M., Effect of ladostigil treatment of aging rats on gene expression in four brain areas associated with regulation of memory. Neuropharmacology 2020, 177, 108229.

21. Reichert, F.; Rotshenker, S., Complement-receptor-3 and scavenger-receptor-AI/II mediated myelin phagocytosis in microglia and macrophages. Neurobiol Dis 2003, 12, (1), 65–72.

22. Gitik, M.; Liraz-Zaltsman, S.; Oldenborg, P. A.; Reichert, F.; Rotshenker, S., Myelin down-regulates myelin phagocytosis by microglia and macrophages through interactions between CD47 on myelin and SIRPalpha (signal regulatory protein-alpha) on phagocytes. J Neuroinflammation 2011, 8, 24.

23. Shamash, S.; Reichert, F.; Rotshenker, S., The cytokine network of Wallerian degeneration: tumor necrosis factor-alpha, interleukin-1alpha, and interleukin-1beta. J Neurosci 2002, 22, (8), 3052–60.

24. He, Y.; Taylor, N.; Fourgeaud, L.; Bhattacharya, A., The role of microglial P2X7: modulation of cell death and cytokine release. J Neuroinflammation 2017, 14, (1), 135.

25. Brown, J.; Pirrung, M.; McCue, L. A., FQC Dashboard: integrates FastQC results into a web-based, interactive, and extensible FASTQ quality control tool. Bioinformatics 2017, 33, (19), 3137–3139.

26. Bolger, A. M.; Lohse, M.; Usadel, B., Trimmomatic: a flexible trimmer for Illumina sequence data. Bioinformatics 2014, 30, (15), 2114–2120.

27. Dobin, A.; Davis, C. A.; Schlesinger, F.; Drenkow, J.; Zaleski, C.; Jha, S.; Batut, P.; Chaisson, M.; Gingeras, T. R., STAR: ultrafast universal RNA-seq aligner. Bioinformatics 2013, 29, (1), 15–21.

28. Robinson, M. D.; McCarthy, D. J.; Smyth, G. K., edgeR: a Bioconductor package for differential expression analysis of digital gene expression data. Bioinformatics 2010, 26, (1), 139–40.

29. Eden, E.; Navon, R.; Steinfeld, I.; Lipson, D.; Yakhini, Z., GOrilla: a tool for discovery and visualization of enriched GO terms in ranked gene lists. BMC Bioinformatics 2009, 10, 48.

30. Szklarczyk, D.; Franceschini, A.; Wyder, S.; Forslund, K.; Heller, D.; Huerta-Cepas, J.; Simonovic, M.; Roth, A.; Santos, A.; Tsafou, K. P., STRING v10: protein–protein interaction networks, integrated over the tree of life. Nucleic acids research 2014, 43, (D1), D447–D452.

31. Pulido-Salgado, M.; Vidal-Taboada, J. M.; Barriga, G. G.; Sola, C.; Saura, J., RNA-Seq transcriptomic profiling of primary murine microglia treated with LPS or LPS + IFNgamma. Sci Rep 2018, 8, (1), 16096.

32. Zhang, J. D.; Wiemann, S., KEGGgraph: a graph approach to KEGG PATHWAY in R and bioconductor. Bioinformatics 2009, 25, (11), 1470–1471.

33. Christofferson, D. E.; Li, Y.; Yuan, J., Control of life-or-death decisions by RIP1 kinase. Annu Rev Physiol 2014, 76, 129–50.

34. Gomez-Nicola, D.; Valle-Argos, B.; Nieto-Sampedro, M., Blockade of IL-15 activity inhibits microglial activation through the NFkappaB, p38, and ERK1/2 pathways, reducing cytokine and chemokine release. Glia 2010, 58, (3), 264–76.

35. Gao, T.; Jernigan, J.; Raza, S. A.; Dammer, E. B.; Xiao, H.; Seyfried, N. T.; Levey, A. I.; Rangaraju, S., Transcriptional regulation of homeostatic and disease-associated-microglial genes by IRF1, LXRbeta, and CEBPalpha. Glia 2019, 67, (10), 1958–1975.

36. Panda, D.; Gjinaj, E.; Bachu, M.; Squire, E.; Novatt, H.; Ozato, K.; Rabin, R. L., IRF1 Maintains Optimal Constitutive Expression of Antiviral Genes and Regulates the Early Antiviral Response. Front Immunol 2019, 10, 1019.

37. He, Y.; Taylor, N.; Yao, X.; Bhattacharya, A., Mouse primary microglia respond differently to LPS and poly(I:C) in vitro. Sci Rep 2021, 11, (1), 10447.

38. Matcovitch-Natan, O.; Winter, D. R.; Giladi, A.; Vargas Aguilar, S.; Spinrad, A.; Sarrazin, S.; Ben-Yehuda, H.; David, E.; Zelada Gonzalez, F.; Perrin, P.; Keren-Shaul, H.; Gury, M.; Lara-Astaiso, D.; Thaiss, C. A.; Cohen, M.; Bahar Halpern, K.; Baruch, K.; Deczkowska, A.; Lorenzo-Vivas, E.; Itzkovitz, S.; Elinav, E.; Sieweke, M. H.; Schwartz, M.; Amit, I., Microglia development follows a stepwise program to regulate brain homeostasis. Science 2016, 353, (6301), aad8670.

39. Mosher, K. I.; Wyss-Coray, T., Microglial dysfunction in brain aging and Alzheimer’s disease. Biochemical pharmacology 2014, 88, (4), 594–604.

40. McInturff, J. E.; Modlin, R. L.; Kim, J., The role of toll-like receptors in the pathogenesis and treatment of dermatological disease. J Invest Dermatol 2005, 125, (1), 1–8.

41. Lynch, M. A., The multifaceted profile of activated microglia. Molecular neurobiology 2009, 40, (2), 139–156.

42. Venegas, C.; Heneka, M. T., Danger-associated molecular patterns in Alzheimer’s disease. J Leukoc Biol 2017, 101, (1), 87–98.

43. Gulke, E.; Gelderblom, M.; Magnus, T., Danger signals in stroke and their role on microglia activation after ischemia. Ther Adv Neurol Disord 2018, 11, 1756286418774254.

44. York, E. M.; Bernier, L. P.; MacVicar, B. A., Microglial modulation of neuronal activity in the healthy brain. Developmental neurobiology 2018, 78, (6), 593–603.

45. Raouf, R.; Chabot-Doré, A.-J.; Ase, A. R.; Blais, D.; Séguéla, P., Differential regulation of microglial P2X4 and P2X7 ATP receptors following LPS-induced activation. Neuropharmacology 2007, 53, (4), 496–504.

46. Parvathenani, L. K.; Tertyshnikova, S.; Greco, C. R.; Roberts, S. B.; Robertson, B.; Posmantur, R., P2X7 mediates superoxide production in primary microglia and is up-regulated in a transgenic mouse model of Alzheimer’s disease. Journal of Biological Chemistry 2003, 278, (15), 13309–13317.

47. Illes, P., P2X7 Receptors Amplify CNS Damage in Neurodegenerative Diseases. Int J Mol Sci 2020, 21, (17).

48. Wong, E. T.; Tergaonkar, V., Roles of NF-κB in health and disease: mechanisms and therapeutic potential. Clinical science 2009, 116, (6), 451–465.

49. Kiyota, T.; Gendelman, H. E.; Weir, R. A.; Higgins, E. E.; Zhang, G.; Jain, M., CCL2 affects β-amyloidosis and progressive neurocognitive dysfunction in a mouse model of Alzheimer’s disease. Neurobiology of aging 2013, 34, (4), 1060–1068.

50. Joly-Amado, A.; Hunter, J.; Quadri, Z.; Zamudio, F.; Rocha-Rangel, P. V.; Chan, D.; Kesarwani, A.; Nash, K.; Lee, D. C.; Morgan, D., CCL2 overexpression in the brain promotes glial activation and accelerates tau pathology in a mouse model of tauopathy. Frontiers in Immunology 2020, 11, 997.

51. Chhor, V.; Le Charpentier, T.; Lebon, S.; Ore, M. V.; Celador, I. L.; Josserand, J.; Degos, V.; Jacotot, E.; Hagberg, H.; Savman, K.; Mallard, C.; Gressens, P.; Fleiss, B., Characterization of phenotype markers and neuronotoxic potential of polarised primary microglia in vitro. Brain Behav Immun 2013, 32, 70–85.

52. Butovsky, O.; Jedrychowski, M. P.; Moore, C. S.; Cialic, R.; Lanser, A. J.; Gabriely, G.; Koeglsperger, T.; Dake, B.; Wu, P. M.; Doykan, C. E.; Fanek, Z.; Liu, L.; Chen, Z.; Rothstein, J. D.; Ransohoff, R. M.; Gygi, S. P.; Antel, J. P.; Weiner, H. L., Identification of a unique TGF-beta-dependent molecular and functional signature in microglia. Nat Neurosci 2014, 17, (1), 131–43.

53. Henn, A.; Lund, S.; Hedtjärn, M.; Schrattenholz, A.; Pörzgen, P.; Leist, M., The suitability of BV2 cells as alternative model system for primary microglia cultures or for animal experiments examining brain inflammation. ALTEX: Alternatives to animal experimentation 2009, 26, (2), 83–94.

54. Mendonca, P.; Taka, E.; Bauer, D.; Cobourne-Duval, M.; Soliman, K. F., The attenuating effects of 1, 2, 3, 4, 6 penta-O-galloyl-β-d-glucose on inflammatory cytokines release from activated BV-2 microglial cells. Journal of neuroimmunology 2017, 305, 9–15.

55. Lively, S.; Schlichter, L. C., Microglia Responses to Pro-inflammatory Stimuli (LPS, IFNgamma+TNFalpha) and Reprogramming by Resolving Cytokines (IL-4, IL-10). Front Cell Neurosci 2018, 12, 215.

56. Hobbs, S.; Reynoso, M.; Geddis, A. V.; Mitrophanov, A. Y.; Matheny, R. W., Jr., LPS-stimulated NF-kappaB p65 dynamic response marks the initiation of TNF expression and transition to IL-10 expression in RAW 264.7 macrophages. Physiol Rep 2018, 6, (21), e13914.

57. Heyen, J. R.; Ye, S.-m.; Finck, B. N.; Johnson, R. W., Interleukin (IL)-10 inhibits IL-6 production in microglia by preventing activation of NF-κB. Molecular Brain Research 2000, 77, (1), 138–147.

58. Thawkar, B. S.; Kaur, G., Inhibitors of NF-κB and P2X7/NLRP3/Caspase 1 pathway in microglia: Novel therapeutic opportunities in neuroinflammation induced early-stage Alzheimer’s disease. Journal of neuroimmunology 2019, 326, 62–74.

59. Shieh, C. H.; Heinrich, A.; Serchov, T.; van Calker, D.; Biber, K., P2X7 dependent, but differentially regulated release of IL 6, CCL2, and TNF α in cultured mouse microglia. Glia 2014, 62, (4), 592–607.

60. Shoham, S.; Linial, M.; Weinstock, M., Age-Induced Spatial Memory Deficits in Rats Are Correlated with Specific Brain Region Alterations in Microglial Morphology and Gene Expression. J Neuroimmune Pharmacol 2019, 14, (2), 251–262.

