## Supplementary material for "Profiles and Dynamics of the Transcriptome of Microglial Cells Reveal their Inflammatory Status": Figures S1-S5

**Supplementary Tables:**

- Table S1: RNA-seq results for bzATP treatment (by TMM)
- Table S2: Gene set of DE genes exposed to bzATP with slow kinetics (source of Table 2)
- Table S3: RNA-seq results for 33 interleukin-related genes (IL-gene set; source of Figure 4C).
- Table S4: RNA-seq results for bzATP/LPS treatment (by TMM)

**Supplementary Figures:**

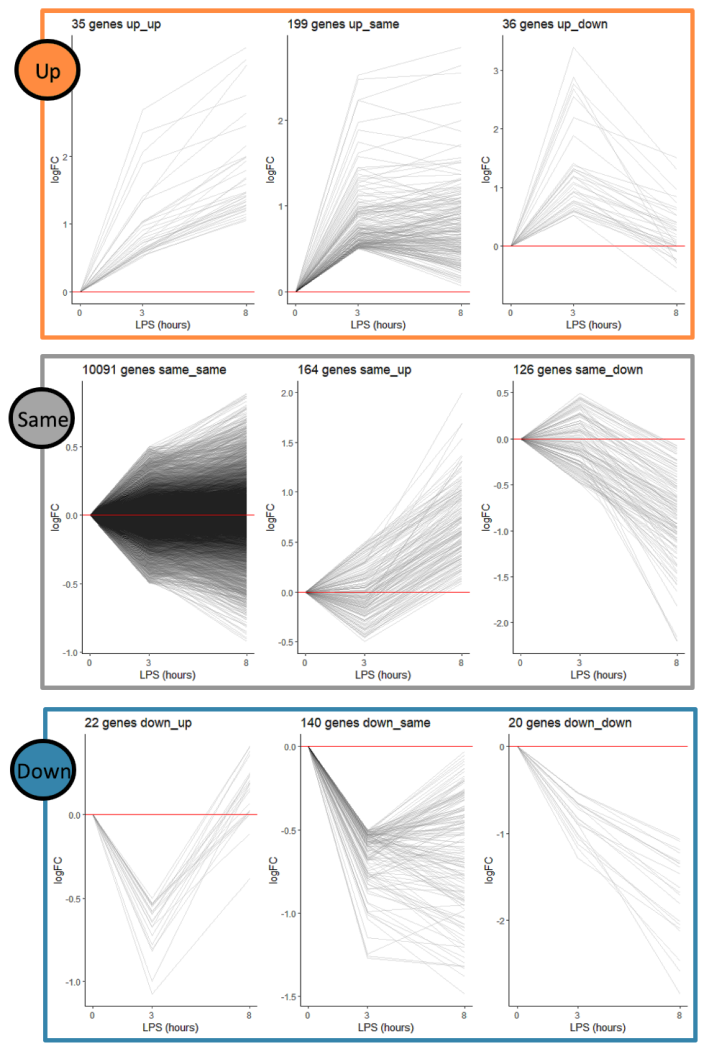

**Figure S1.** Gene expression trends on naïve, not treated microglial cells and following activation with bzATP for 3 hrs and 8 hrs. Each differentially expressed gene is shown as a gray line connecting the hours from the exposure. The y-axis shows the log (base 2) fold change (log FC). The horizontal red line indicates a baseline with no change in expression. Note that the scale of the y-axis changes according to the range of the genes in the clustered group. The number of genes associated with each cluster group (Marked as Up, Same and Down) are indicated. Analysis is based on data in **Supplemental Table S1**.

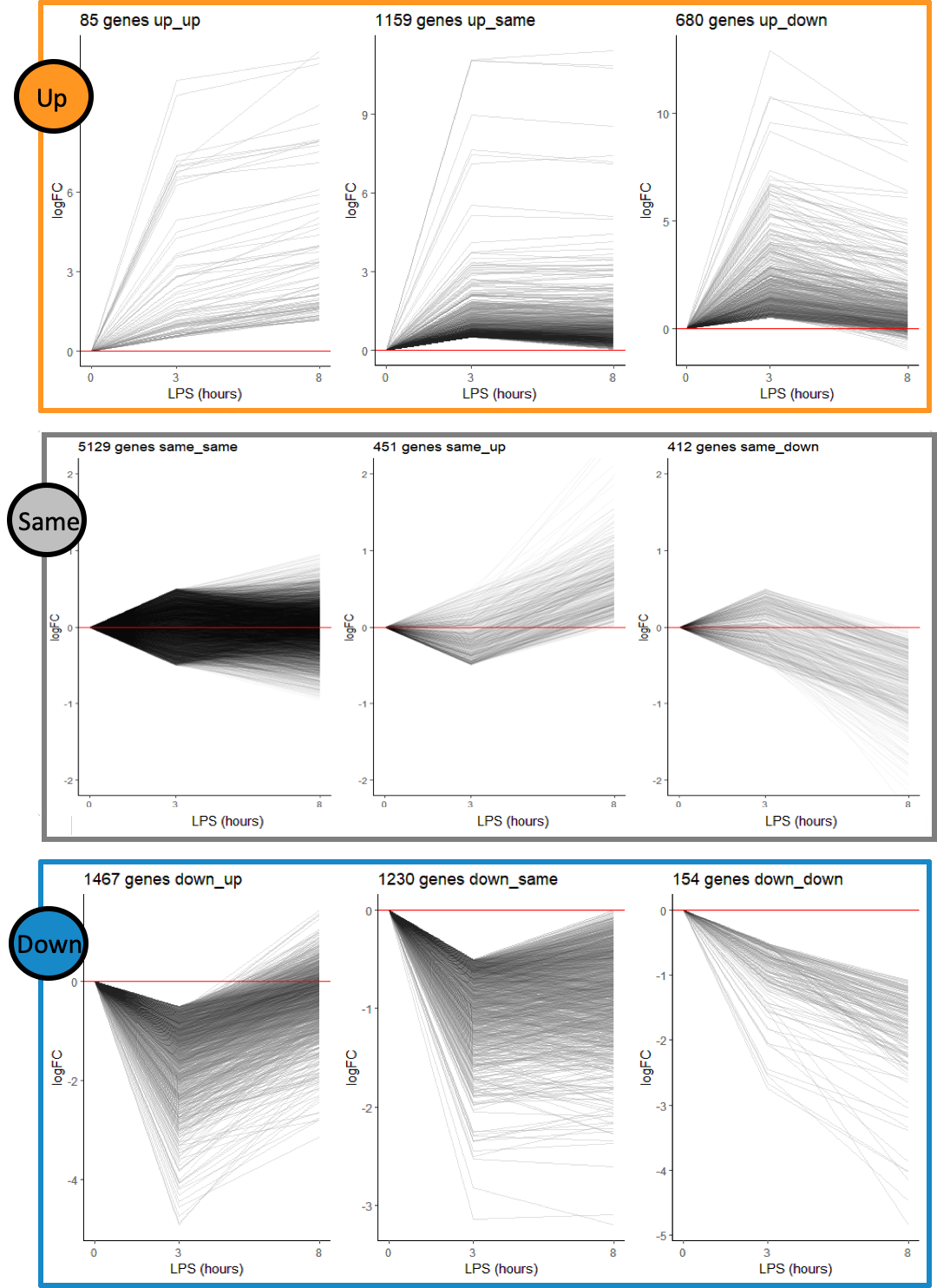

**Figure S2.** Gene expression trends on naïve, not treated microglial cells and following activation with bzATP/LPS for 3 hrs and 8 hrs. Each differentially expressed gene is shown as a gray line connecting the hours from the exposure. The y-axis shows the log (base 2) fold change (log FC). The horizontal red line indicates a baseline with no change in expression. Note that the scale of the y-axis changes according to the range of the genes in the clustered group. The number of genes associated with each cluster group (Marked as Up, Same and Down) are indicated. Analysis is based on data in **Supplemental Table S4**.

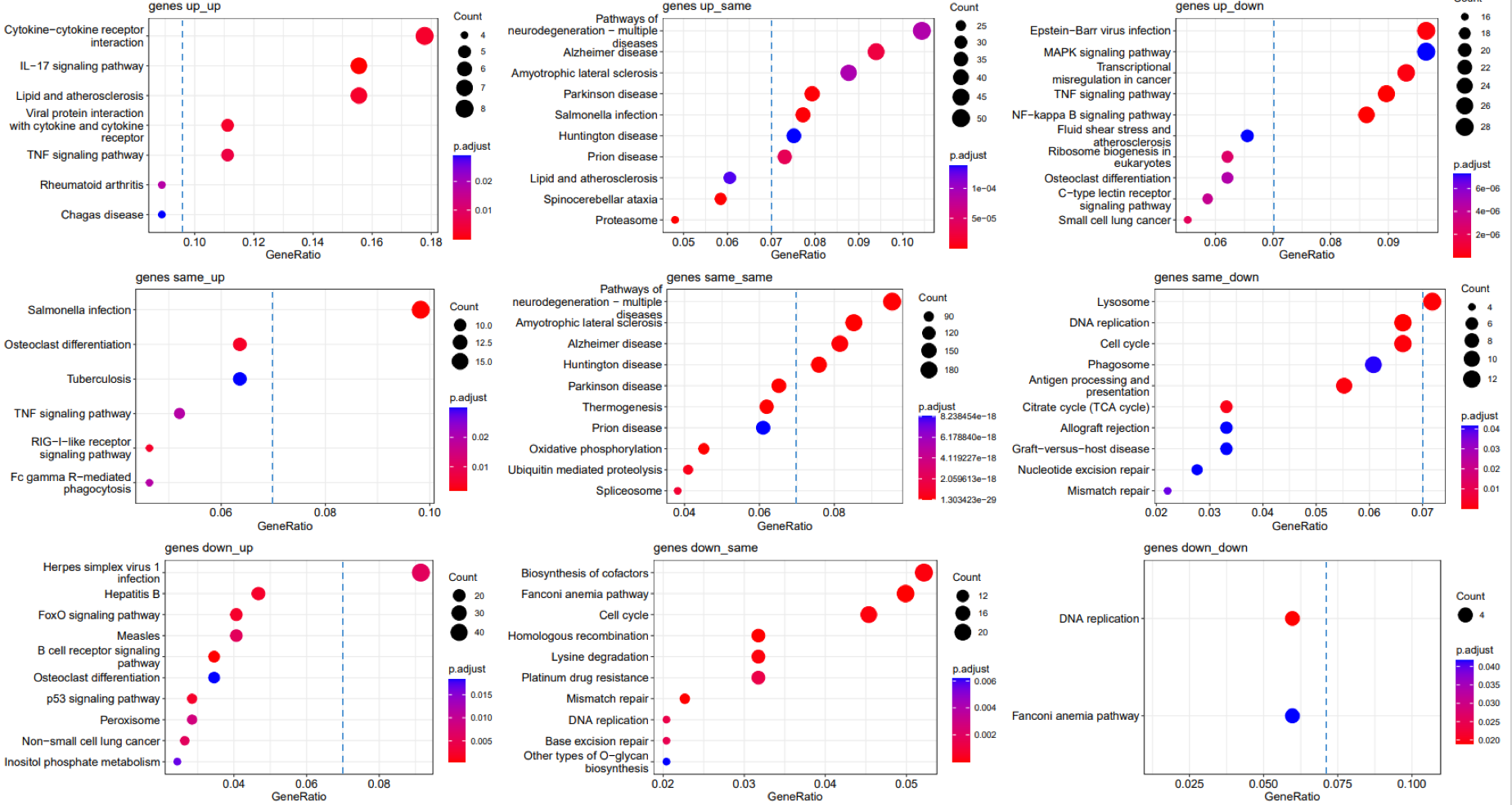

**Figure S3:** Enrichment analysis for all 9 differentially expressed (DE) gene clusters followed bzATP/LTP induction. Enrichment analysis for all DE genes clusters. Top, middle and bottom panels are associated with Up, Same and Down set (3 hrs). Annotations are according to KEGG pathway resource. Statistical enrichment is depicted by p-adjust and significance is bounded to FDR <0.05. The size of the dots captures the number of proteins. Note that the enrichment significance scale, the number of genes associated with each enriched pathway, and the gene ratio (i.e., fraction of genes in the cluster that are included in the specific pathway) differ among gene cluster. A vertical dashed line marks Gene ratio = 0.7 for better visual alignment.

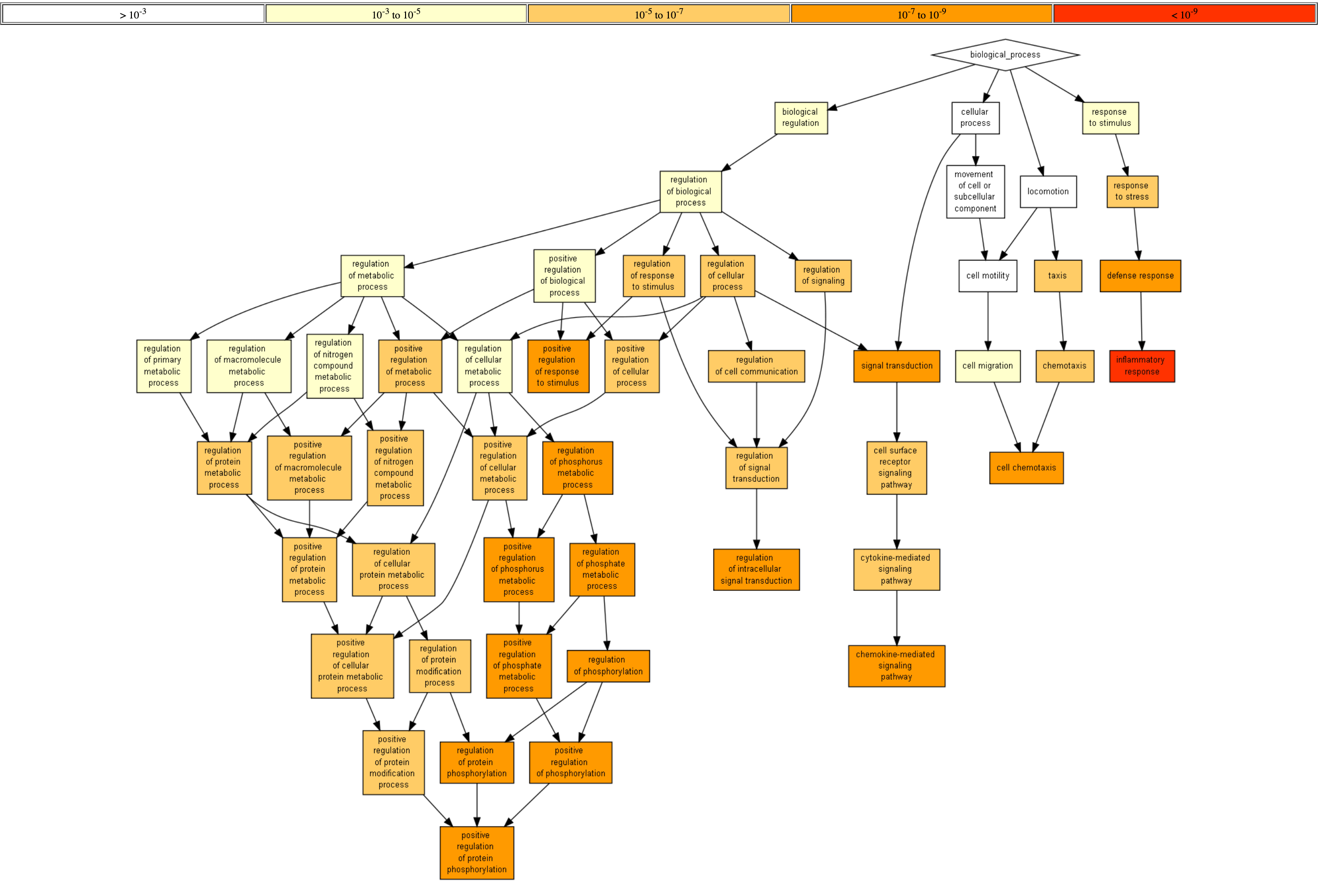

**Figure S4.** Enrichment analysis for gene ontology (GO) of upregulated DE genes. An enrichment test was applied on 585 statistically significant DE genes that were strongly induced (>1.5-fold increase in expression relative to that of non-treated cells, and average expression level >10 TMM). The test was based on hypergeometric analysis, followed by multiple testing corrections, (GOrilla statistical tool). Results shown are for biological process. Only GO terms with p-value <1.0e-07 were included in the analysis. The color scale (white to red) with the associated range of p-values is shown (Top).

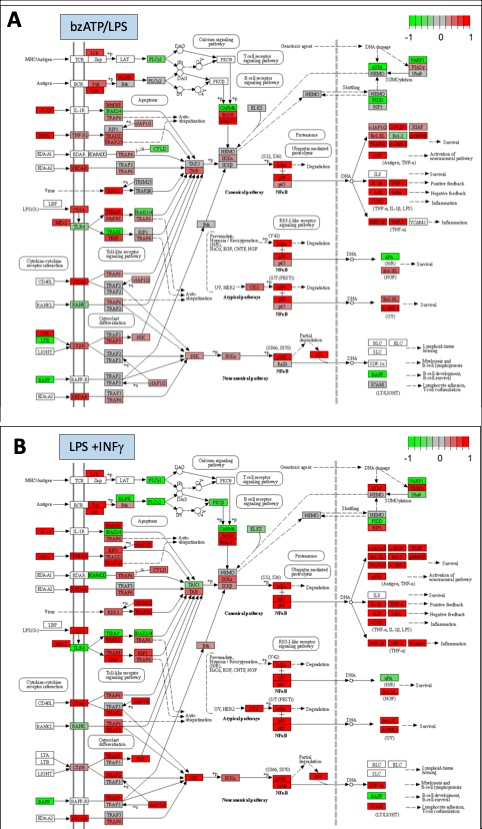

**Figure S5.** DE gene pattern on the KEGG pathway map of NF-κB signaling. The top panel indicates the genes that are downregulated (green) upregulated (red), unchanged (Gray), and undetected (white). The DE genes in cells exposed to bzATP/LPS (8 hrs, Top), and LPS+IFNγ for 6 hrs (Bottom) are shown.
